# Does targeted memory reactivation during NREM sleep require complementary REM sleep for memory consolidation?

**DOI:** 10.1101/2023.05.08.539563

**Authors:** Rebeca Sifuentes Ortega, Péter Simor, Philippe Peigneux

## Abstract

Presentation of learning-related cues during NREM sleep has been shown to improve memory consolidation. Past studies suggest that REM sleep may contribute to the beneficial effect of reactivating memories during NREM sleep, but the relationship between REM sleep and induced reactivations in NREM remains unclear. We investigated whether a naturally ensuing episode of REM sleep is necessary for prior NREM targeted memory reactivation (TMR) to exert a beneficial effect on memory consolidation. Nineteen participants learned the association between prior or non-prior known objects and their names (pseudowords) in a within-subject multiple session experiment, in which TMR was subsequently performed either before (Pre-REM) or after (Post-REM) the final REM sleep episode of the night. While word-picture association recall measures did not differ between TMR conditions, we found better name recognition based on confidence ratings for words reactivated during Pre-REM TMR in contrast with associations cued during Post-REM TMR. In addition, we found distinct associations between cue-evoked sigma activity, subsequent REM theta power and TMR memory benefits which were contingent upon the level of relatedness with prior knowledge for the learned material. Although TMR may be less effective during the second half of the night, our findings suggest an interplay between NREM and REM sleep oscillatory activity for memory reactivation and consolidation processes.

## Introduction

Targeted memory reactivation (TMR) in sleep, or the external reactivation of memory traces through the presentation of learning-related cues, has proven useful to understand post-learning memory consolidation processes taking place during sleep. A growing body of evidence has shown that applying TMR, particularly during non-rapid eye movement sleep (NREM), can result in improved performance in various memory domains (Hu et al. 2020). This improvement is believed to result from the coordination of oscillatory phenomena characterising NREM sleep. Namely, hippocampal sharp-wave ripples (SWR), associated to memory reactivations (Buzsáki 2015), are nested within thalamic spindles (Siapas and Wilson 1998; Staresina et al. 2015) which are coupled with the up-states of neocortical slow oscillations (Sirota et al. 2003; Latchoumane et al. 2017). This integrated mechanism would support the consolidation and integration of hippocampus-dependent memory into neocortical long-term stores (Buzsáki 1996; Diekelmann and Born 2010; Klinzing et al. 2019).

Recently acquired memories are thought to undergo continuous transformations throughout post-encoding sleep, characterized by alternating NREM-REM cycles. Several accounts assign complementary roles to these sleep stages in the consolidation and transformation of memories (Giuditta et al. 1995; Ambrosini and Giuditta 2001; Diekelmann and Born 2010). In the framework of the active systems consolidation model, recent memories are first reorganized by coordinated oscillatory activities in NREM slow wave sleep (SWS), then strengthened at the synaptic level during subsequent REM sleep (Diekelmann and Born 2010; Klinzing et al. 2019). Alternatively, the sequential hypothesis account proposes that it is the physiologically ordered NREM-REM sleep sequence that plays a key role for memory consolidation (Giuditta et al. 1995). In SWS, non-adaptive memory traces would be weakened or eliminated, leading to a relative strengthening of the adaptive traces, and then these traces would be consolidated and integrated into long-term stores during a following episode of REM sleep. Evidence supporting the sequential hypothesis in humans showed that ordered NREM-REM sequences are relevant for declarative memory (Ficca et al. 2000) and visual perceptual skill learning (Strauss et al. 2022). Even though these models consider that consolidation processes continue throughout both sleep stages, little is known about the role played by REM sleep after TMR-triggered memory reactivations during prior NREM sleep.

In two novel word learning studies, REM sleep was found to exert an influence on the outcome of prior NREM-TMR. Batterink et al. (2017) found that NREM sleep TMR during a nap did not benefit memory performance for cued words (i.e., pseudowords associated with pictures of known objects at learning) at the group level, at variance with previous studies disclosing an advantage for cued foreign language word pairs (Schreiner and Rasch 2015; Schreiner et al. 2015), and pseudowords associated with familiar objects (Groch et al. 2017). Notwithstanding, Batterink et al. (2017) observed the expected TMR benefit in participants who obtained the greatest amounts of REM sleep, whereas those with little to no REM sleep paradoxically exhibited worse memory performance for cued than uncued associations. Similarly, Tamminen et al. (2017) found no NREM-TMR effect on the retrieval of novel words (e.g., *cathedruke* derived from base word *cathedral*), but found that the change in lexical competition (a measure of lexical integration) was associated with the amount of time spent in REM sleep during the afternoon nap. Words cued during NREM sleep benefited from subsequent REM sleep and exhibited increased lexical competition, whereas such relationship was not observed for uncued words. These results suggest that REM sleep following the reactivation of novel vocabulary may be particularly beneficial because these type of associations require further integration within pre-existing knowledge networks (Batterink et al. 2017; Tamminen et al. 2017), as opposed to simpler associations that could still benefit from TMR in the absence of REM sleep (Hu et al. 2020).

Another line of evidence shows that offline periods filled with sleep facilitate novel word integration (Gais et al. 2006; Dumay and Gaskell 2007; Tamminen et al. 2010; Schimke et al. 2021). In particular, sleep spindles and SWS have been associated with overnight improvements in novel word recognition and lexical competition (Tamminen et al. 2010). Moreover, NREM-TMR studies showing a benefit in the recall of foreign language word pairs (Schreiner and Rasch 2015; Schreiner et al. 2015) or pseudoword-picture associations (Groch et al. 2017) also featured increased theta and sigma activity (related to sleep spindles) after the presentation of learning-related cues. Notably in the latter study, the increase in cue-evoked theta and sigma activity, as well as the TMR memory benefit, were selective for cued pseudowords previously associated with pictures of familiar objects, but not for those paired with unfamiliar objects (Groch et al. 2017). Furthermore, the increases in sigma and theta oscillatory activity predicted a TMR benefit only for prior known associations, supporting the notion that prior knowledge is an essential element for sleep-related consolidation processes to unfold (Durrant et al. 2015; Hennies et al. 2016; Groch et al. 2017; but see Ashton et al. 2022; Cordi et al. 2023 for a challenging view). It is worth noticing that in these novel vocabulary TMR studies, cueing was performed during the first half of the night, allowing participants to spend at least some time in REM sleep (Schreiner and Rasch 2015; Schreiner et al. 2015; Groch et al. 2017). Still, it remains to be systematically tested whether a period of REM sleep *after* NREM-TMR is relevant for strengthening and integrating new memories, and whether this depends on the level of relatedness of the newly learned material with prior knowledge.

In this context, we aimed at investigating whether NREM-cued memories benefit from a subsequent episode of REM sleep, and if this potential benefit is contingent upon prior knowledge. To this end, we performed a within-subject study in which TMR took place either before or after a final episode of REM sleep, while controlling for sleep duration. In line with the Sequential Hypothesis (Giuditta et al. 1995; Ambrosini and Giuditta 2001) and studies showing an influence of REM sleep on the beneficial effects of NREM-TMR for highly integrative stimuli such as novel words (Batterink et al. 2017; Tamminen et al. 2017), we hypothesized that memory performance would benefit from NREM cueing only when followed by a period of REM sleep (Pre-REM TMR condition). In particular, we predicted that cued associations that involve prior knowledge would benefit to a greater extent from TMR, given their consistent links with pre-existing knowledge networks, and from subsequent REM sleep proposed to integrate previously reactivated memories (Batterink et al. 2017; Tamminen et al. 2017). On the other hand, we hypothesized that cueing memories during NREM sleep without a subsequent period of REM sleep (Post-REM TMR condition) may lead to the destabilization of the memory traces, and consequently a decrease in memory performance for cued associations. This decrease would be especially prominent for associations involving less connections with prior knowledge, since these would benefit to a less extent from the reactivation processes promoting integration into pre-existing networks. Additionally, given previous findings on the increase in sigma activity and its association with TMR memory benefits, we investigated the oscillatory activity related to cue presentation in both TMR conditions, and the potential role of REM theta activity following a period of NREM TMR. We focused on REM sleep theta activity given its role in memory consolidation processes (Boyce et al. 2016, 2017) and its observed role in modulating hippocampal SWR activity, likely associated to memory reactivations, over the course of NREM-REM cycles (Grosmark et al. 2012).

## Materials and Methods

### Participants

Twenty-two healthy participants (10 males, mean age 24.6 ± 3.5 years, range 20-33) gave written informed consent to take part in this study approved by the ULB-Erasme Hospital Ethics Committee (Ref. P2019/332). They were French-native speakers, with no history of neurological or psychiatric disorders, no signs of depressive symptoms (short form of the Beck Depression Inventory BDI-13; Beck and Beck 1972; mean score = 1.2 ± 1.9), free of medication, with reported good sleep quality (Pittsburgh Sleep Quality Index PSQI; Buysse et al., 1989; mean score = 2.6 ± 1.2), and neutral to moderate chronotype (Morningness-Eveningness Questionnaire; Horne and Östberg, 1976; mean score = 56.5 ± 5.2). Three participants were excluded from the analyses due to poor sleep during the experimental nights and REM sleep intrusion in the Post-REM cueing condition (n= 1), dropping out of the study (n=1), or poor performance at the immediate recall test (i.e., scoring 3 times below the median absolute deviation; n = 1). Data are thus analysed on 19 participants (8 males, mean age 24.4 ± 3.6 years). All participants were instructed to refrain from consuming alcohol, taking caffeine or stimulant drinks after noon, or napping on the experimental days. After completion of the experiment, they received a monetary compensation.

### Materials

One hundred and fifty 2D pictures of known and unknown objects, adapted from a previous study (Urbain et al. 2013) and pre-existing databases (Kroll and Potter 1984; Rossion and Pourtois 2004), were assigned to three different lists consisting each of 25 prior known and 25 non-prior known objects. Each picture (prior- and non-prior known) was paired with a pseudoword and a “magical” function of the object (Fig. 1). Pseudowords were generated using the Lexique toolbox (New et al. 2001) according to French language rules, and balanced across lists in syllable number (mean syllable length = 2.46, range 1-3) and word duration (mean = 794.9 ± 172.9 ms). Picture definitions were 5 to 13 words long. Pseudowords and definitions were recorded by a female French native speaker. Stimuli were presented using PsychoPy 3 (Peirce et al. 2019).

**Figure 1.**
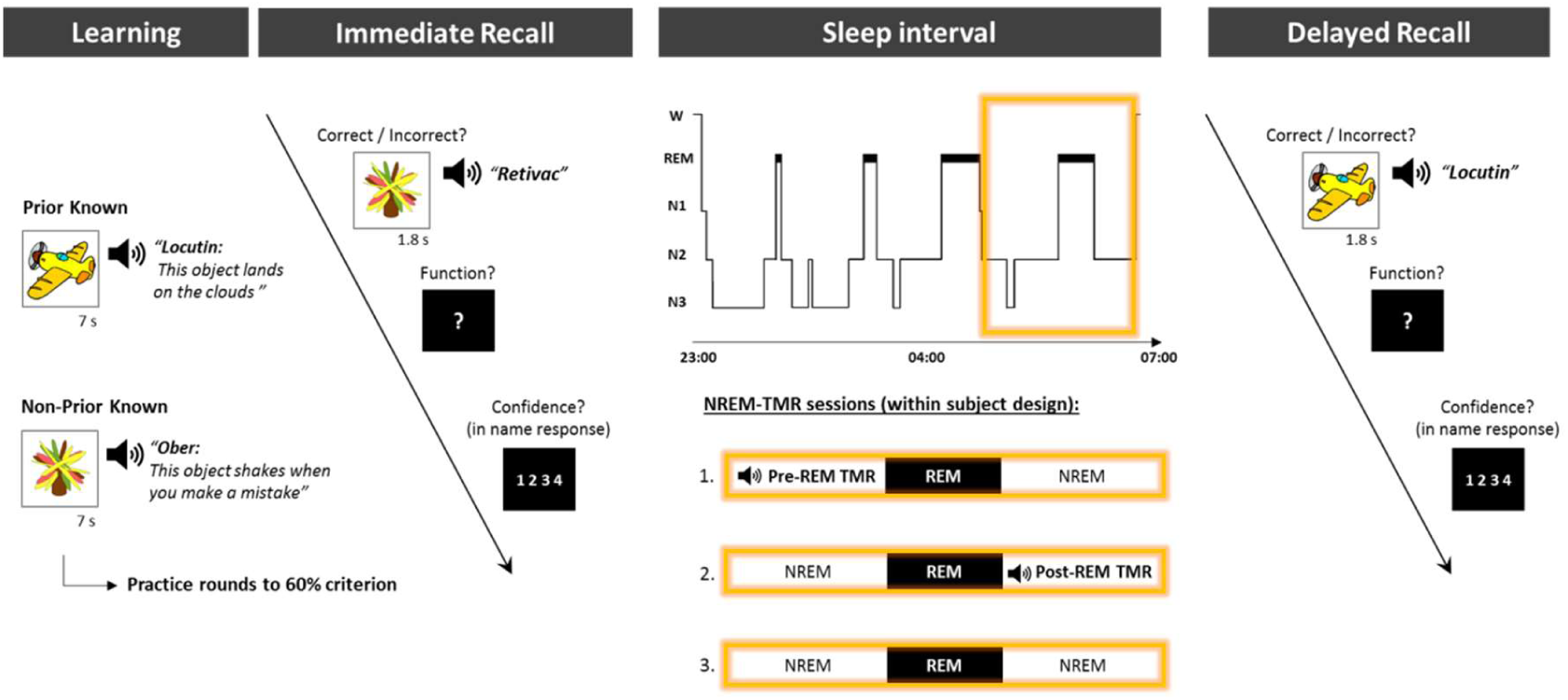
Experimental design. After a habituation night, participants took part in 3 experimental sessions. In each session, three different lists of word-picture associations were presented. During the encoding phase, participants learned 50 new pseudowords associated to a prior known or non-prior known object and their “magical” function. After reaching a 60% learning criterion in subsequent practice rounds, immediate recall was tested. In this phase, participants were asked to rate their confidence on the correctness of their name recognition responses. During the sleep interval, sleep signals were monitored and TMR was performed during NREM sleep either before a REM sleep episode (Pre-REM cueing) or after a REM sleep episode (Post-REM cueing), or participants slept undisturbed (Control). After the sleep interval, a delayed recall test took place. REM; rapid-eye movement sleep, NREM; non-rapid eye movement sleep.

### General procedure

In total, each participant took part in 4 experimental sessions, each separated by at least one week. The first session was the habituation night to familiarize participants with the polysomnography (PSG) setup and the sleep lab environment. For the three other sessions, the experimental night started at 20:30 with the setup of the electrodes, followed by a learning and immediate recall phase, then nocturnal sleep in one of the 3 experimental conditions (see below), and delayed recall in the morning in all cases (Fig.1). At each experimental night, participants learned a different list of 50 pseudoword-picture associations. List order was counterbalanced across conditions. According to the experimental condition, TMR of previously learned words was performed during the second half of the night during NREM sleep either (a) before the last expected REM episode in the night (Pre-REM TMR) or (b) after the last REM sleep episode (Post-REM TMR), or (c) no TMR was performed — i.e., sleep control. The order of the experimental conditions was counterbalanced across participants. Thirty minutes after participants were woken up, the delayed recall test was administered.

### Pseudoword-picture association learning

In the initial learning phase, participants had to memorize the association between 50 pictures of prior and non-prior known objects, their names consisting of pseudowords, and their “magical” functions. Each object picture was displayed on a computer screen for 7 s together with its name and function which were presented auditorily through earphones. After the presentation of each association, a question mark replaced the object picture, and the participant was asked to repeat verbally the name and function of the object. Associations were presented in blocks of five, after which participants were presented again with each of the last five objects and their names (1.8 s) and were asked to recall once again the complete association when the question mark appeared. After completing the learning procedure with all 50 associations, a practice round followed.

For the practice round, each object picture was presented (1.8 s) either with its corresponding name or an incorrect one (targets and lures respectively, presented 50% of the time for each type of knowledge). Next, a question mark appeared on the screen and participants had to judge whether the name corresponded or not to the object — i.e., by choosing between “correct” or “incorrect”— and to recall the object function. Functions, scored online by the experimenter, were scored as incorrect only when partially retrieved (e.g., *“This object shakes*” instead of *“This object shakes when you make a mistake”*), or when the real definition of a familiar object was given instead of the magical function (e.g., *“This object flies”* instead of *“This object lands on the clouds”;* see Fig.1). Participants had to reach 60% retrieval accuracy for full associations (i.e., correctly retrieved name and function). If the criterion was not reached, participants went through a new learning and practice round including only incorrectly remembered associations. Once participants reached the 60% criterion of the total of associations (that is, including the ones previously recalled correctly), they were administered an immediate recall test as explained below.

### Immediate and delayed recall tests

Each of the 50 object pictures was presented with an associated name, and participants had then to judge whether it was “correct” or “incorrect” and recall its function. Immediately after each object presentation, they rated their confidence in their name recognition response on a scale ranging from 1 “not sure at all” to 4 “completely sure” on a keyboard (see Fig. 1).

### Targeted Memory Reactivation

After completion of the immediate recall test, participants went to sleep. In both the Pre-REM and Post-REM TMR conditions, half of the prior known and non-prior known associations fully remembered at the immediate recall test were randomly assigned either to a “cueing” or “no cueing” word subset using an algorithm implemented in Matlab R2017b (https://www.mathworks.com). Only fully remembered associations at immediate recall were chosen in order to compare the effects of TMR on post-sleep retention of cued versus uncued associations. In other words, the subset of uncued words served as a reference to compare the effects of TMR on the next day’s memory performance at delayed recall. The word list from the cueing subset was repeatedly played during the selected NREM sleep episode at ∼40 dB via insert earphones. The inter-cue interval was 9 s. TMR was performed when stable NREM sleep was observed during online monitoring of the PSG signals, and paused whenever signs of arousal appeared. In the Pre-REM TMR condition, cueing was performed within the NREM period preceding the last REM episode expected in the night, starting approximately during the 4^th^ sleep cycle. After cueing, participants were allowed to transition to the following REM episode and were awoken at the end of the subsequent NREM episode. In the Post-REM TMR condition, NREM cueing started approximately on the 5^th^ sleep cycle, right after the last REM episode of the night was over. For this condition, participants were awoken towards the end of the cued NREM episode, hence avoiding the transition to a subsequent REM episode.

### Polysomnography

Continuous EEG recordings were conducted using an Active-Two Biosemi system (Biosemi, The Netherlands) with active Ag-AgCl electrodes at the 19 scalp locations of the 10-20 international system, as well as left and right mastoids, and standard electrooculography (EOG) and electromyography (EMG) derivations (Berry et al. 2017). Signals were referenced to the Common Mode Sense (CMS) electrode and digitized at 2048 Hz sampling rate. For EEG data pre-processing (for offline scoring and EEG analyses), signals were exported and downsampled to 512 Hz, and EEG channels were re-referenced to averaged mastoids (M1 and M2). EEG and EOG channels were high-passed filtered at 0.3 Hz and low-passed filtered at 45 Hz, and EMG was high- and low-passed filtered at 10 and 50 Hz respectively. Offline sleep scoring was performed on 30-s epochs according to standard criteria (Berry et al. 2017) by an experienced researcher blind to stimulation onsets.

### Behavioural Data Analysis

#### Overall change in memory performance

We first investigated overall changes in memory performance by comparing the total number of fully remembered associations, the number of remembered functions, and the recognition accuracy for object names. Fully remembered associations were defined as correctly retrieving both the name and function associated to an object, whereas name recognition accuracy was assessed with the *d’* sensitivity index, calculated as: z(*p*[hit]) - z(*p*[false alarms]) according to signal detection theory (Stanislaw and Todorov 1999). To avoid undefined values of *d’* due to extreme proportions of hits and false alarms, we applied the log-linear rule (Hautus 1995) before calculating response proportions. The log-linear approach consists in adding 0.5 to the total number of hits and false alarms and adding 1 to the number of trials before calculating each proportion of responses. Overall change in memory performance was compared across the three experimental conditions (Pre-REM, Post-REM, and Control), recall sessions (Immediate vs. Delayed), and type of knowledge involved in the learned associations (Prior vs. Non-Prior) by means of repeated-measures ANOVAs.

#### Cueing effects on memory performance

In a second step we investigated the cueing effects of TMR by focusing on the relative change in memory performance measures within the subset of fully remembered associations at immediate recall. In the TMR conditions, half of the correctly remembered words were randomly assigned to a “cueing” condition, while the other half was allocated to an “uncued” subset of words. In this way, performance was matched between cued and uncued conditions prior to sleep, enabling us to investigate whether TMR elicited any change in memory retention after sleep. Changes in memory for full associations, functions, and name recognition were investigated across TMR conditions (Pre-REM vs. Post-REM), cueing subsets (Cued vs. Uncued) and type of knowledge (Prior vs. Non-prior known) by means of repeated-measures ANOVAs.

#### Cueing effects on name recognition based on confidence ratings

We further assessed the participants’ ability to discriminate correct from incorrect object names after TMR by computing Receiver Operating Characteristic (ROC) curves for each individual and condition at the delayed recall test by using the confidence ratings provided after each recognition trial (Weidemann and Kahana 2016). Binary responses (“correct” / “incorrect”) to target and lure trials were classified according to confidence ratings in an 8-bin scale ranging from “correct” responses with the highest level of confidence (4) to “incorrect” responses with the highest level of confidence (4). Hit and false alarm proportions were calculated at each bin. ROC curves were constructed by plotting the cumulative hit rate (i.e., “correct” responses to targets, and “incorrect” responses to lures) against the false alarm rate (i.e., “incorrect” responses to targets and “correct” responses to lures) for each condition. We next calculated the area under the curve (AUC) for each individual ROC curve by using the trapezoidal rule in order to compare name discrimination across TMR, cueing and prior knowledge conditions by means of a repeated-measures ANOVA.

All behavioural data were analysed in JASP (JASP Team 2023, version 0.14.1). Repeated-measures analysis of variance (ANOVA) were performed as indicated above, post-hoc t-tests were two-sided. Significance level was set at *p* < 0.05 and effect sizes are reported as partial eta squared or Cohen’s d where appropriate.

### EEG Data Analysis

#### Time-frequency analysis of cue-related responses

Pre-processing and analysis of EEG data were performed with the Fieldtrip toolbox (Oostenveld et al. 2011). To investigate the changes in spectral power over time in response to cue presentation (prior and non-prior known words) during both TMR conditions (Pre-REM and Post-REM), we computed time-frequency representations of power (TFRs). To this end, pre-processed continuous signals from sleep (see Polysomnography section above) were segmented into epochs from −3 to 6 s relative to cue onset, which were next sorted according to the type of cue (prior known or non-prior known), and subsequent memory performance (remembered or forgotten at delayed recall). After visual inspection of trials for artefact removal, TFRs were computed by implementing a Fast Fourier Transform analysis on a Hanning window of 2 s duration sliding in increments of 50 ms over trials of −3 to 6 s window length with the ‘mtmconvol’ function from Fieldtrip. Frequencies of interest ranged from 0.5-25 Hz, and a relative baseline correction was performed on the TFRs with respect to the pre-stimulus onset period between −2 to −1 s. Next, TFRs from subsequently remembered cued associations were compared across TMR and type of knowledge conditions. We focused our analysis on subsequently remembered associations given previous findings showing a significant increase in cue-evoked theta and sigma power for later-remembered associations which predicted TMR memory benefits (Groch et al. 2017). A comparison with subsequently forgotten associations was not possible given the few trials available in this latter condition (e.g., ∼80% of cued associations were remembered at delayed recall, see Fig. 3). Power changes in the TFRs were statistically tested with non-parametric cluster-based permutation testing as implemented in Fieldtrip (Maris and Oostenveld 2007).

We further computed Pearson correlations between mean power changes found in the TFR analysis over grouped electrodes according to the following locations: frontopolar (Fp1, Fp2), frontal (F3, Fz, F4, F7, F8), central (C3, Cz, C4), temporal (T3, T4, T5, T6), parietal (P3, Pz, P4) and occipital (O1, O2), and the TMR benefit index for prior known and non-prior known associations. The TMR benefit index was calculated as the relative change in memory performance (from immediate to delayed recall) for fully remembered cued associations minus the same relative change for uncued ones (e.g., Prior Known TMR benefit = %change cued prior known - %change uncued prior known). We chose to calculate the TMR benefit index on fully remembered associations since this is the outcome upon which cued and uncued word subsets were assigned after immediate recall testing to measure cueing effects.

#### Spectral power analysis of REM sleep after TMR

We investigated the relation between theta power (4-8 Hz) in the subsequent REM sleep period and the Pre-REM TMR benefit index for both prior and non-prior known associations. Sleep epochs of the subsequent REM sleep episode in the Pre-REM TMR condition were segmented into consecutive 5-sec epochs with a 50% overlap and tapered with a Hanning window prior to spectral analysis. Power spectra were obtained by means of the Fast Fourier transform at all electrodes and averaged according to the six electrode locations described above. Correlations between REM theta power and the TMR benefit index were calculated. For all correlation analyses, the false discovery rate (FDR) correction was applied for multiple comparisons (Benjamini and Hochberg 1995). Differences between correlations were tested using Steiger’s Z-test as implemented in the *cocor* package (Diedenhofen and Musch 2015) in R (R Core Team, 2019). An overall threshold of *p* < 0.05 was used for statistical significance across EEG analyses.

## Results

### Sleep and cueing parameters

Total sleep time, wake after sleep onset (WASO), and sleep stage duration did not differ significantly between control and Pre- and Post-REM TMR conditions, except for stages 2 and 3 of NREM sleep, where stage 2 duration was longer in both TMR conditions as compared to the control night, whereas stage 3 duration was longer in the control night as compared to both TMR conditions probably due to the TMR intervention lightening NREM sleep (Table 1). By design, REM duration after NREM-TMR significantly differed between the Pre-REM (mean duration = 30.29 min ± 12.9 min) and Post-REM (mean duration = 0.18 min ± 0.3) TMR conditions (t _18_ = 10.09, p < 0.0001, d = 2.34). A closer analysis of the NREM episode during which stimulation was delivered evidenced a significantly longer duration in Pre-REM TMR (mean = 72.53 min ± 17.02) than in Post-REM TMR (mean = 52.74 min ± 16.89; t _18_ = 3.28, p = 0.004, d = 0.75). Proportionally, the amount of NREM stage 1 (Pre-REM = 7.78% ± 5.66; Post-REM = 6.01% ± 7.27), stage 2 (Pre-REM = 80.44% ± 8.63; Post-REM = 85.76% ± 15.31), and stage 3 (Pre-REM = 2.60% ± 5.91; Post-REM = 3.96% ± 12.16), did not differ significantly between NREM stimulation periods (*ps* > 0.19). Conversely, the proportion of wake during the NREM stimulation episode was significantly higher in Pre-REM TMR (mean = 9.18% ± 4.73) than in Post-REM TMR (mean = 4.27% ± 4.92; t _18_ = 2.88, p = 0.01, d = 0.66).

**Table 1.** Sleep parameters.

| Duration (min) | Control | Pre-REM | Post-REM | *F* | *p* | *Control vs. Pre-REM* | *Control vs. Post-REM* | *Pre-REM vs. Post-REM* |
| --- | --- | --- | --- | --- | --- | --- | --- | --- |
| N1 | 18.2 ± 2.5 | 14.8 ± 1.3 | 16.0 ± 2.4 | 1.06 | 0.36 |  |  |  |
| N2 | 203.6 ± 8.2 | 232.1 ± 7.1 | 241.9 ± 7.4 | 9.23 | 0.001* | *t* = -3.08, *p* < 0.01* | *t* = -4.14, *p* < 0.001* | *t* = 1.06, *p* = 0.3 |
| N3 | 119.6 ± 7.5 | 96.0 ± 6.3 | 94.7 ± 8.5 | 12.72 | 0.0001* | *t* = 4.24, *p* < 0.001* | *t* = 4.48, *p* < 0.001* | *t* = -0.24, *p* = 0.8 |
| REM | 94.5 ± 6.4 | 96.3 ± 6.4 | 91.6 ± 5.0 | 0.23 | 0.79 |  |  |  |
| WASO | 31.16 ± 6.7 | 19.18 ± 1.2 | 18.90 ± 2.5 | 2.8 | 0.1 |  |  |  |
| TST | 435.97 ± 14.9 | 443.16 ± 9.5 | 444.37 ± 8.2 | 0.27 | 0.77 |  |  |  |

Sleep architecture parameters (mean ± SEM). N1, N2, N3: NREM sleep stages 1, 2 and 3; REM, rapid eye movement sleep; WASO, wake after sleep onset; TST, total sleep time. \**P* < .05.

As for the cueing material for TMR, the total number of words selected for cueing (Pre-REM TMR = 17.79 ± 2.8; Post-REM TMR = 17.42 ± 2.3, out of 50 associations), and the total of repetitions per word during TMR (Pre-REM TMR = 5.7 ± 2.1; Post-REM TMR = 7.2 ± 3.9) did not significantly differ between TMR conditions (*ps* > 0.17).

### Overall change in memory performance

Recall progress in the pseudoword-picture association task during practice rounds before immediate recall testing is described in the Supplementary Information (Figs. S1). Comparison of overall change in memory performance for fully remembered associations across the three experimental conditions (Pre-REM, Post-REM, and Control) with factors Recall (Immediate vs. Delayed) and Knowledge (Prior vs, Non-Prior) revealed a significant main effect of knowledge (F_1,18_ = 44.12, p < 0.0001, η_p_^2^ = 0.71) with better memory performance for prior known associations (Fig. 2). There were no main effects of recall (p = 0.12, η_p_^2^ = 0.12), TMR condition (p = 0.27, η_p_^2^ = 0.07), nor significant interactions (*ps* > 0.12). A similar pattern of results emerged for overall function recall, with a significant main effect of knowledge (F_1,18_ = 23.32, p = 0.0001, η_p_^2^ = 0.56), and non-significant main effects of recall (p = 0.25, η_p_^2^ = 0.07), TMR experimental condition (p = 0.65, η_p_^2^ = 0.01), nor interactions between factors (*ps* > 0.08) (Fig. 2). Overall accuracy in name recognition was again better for prior known as compared to non-prior known associations (F_1,18_ = 22.72, p = 0.0002, η_p_^2^ = 0.56), with no main effects of recall (p = 0.23, η_p_^2^ = 0.08), condition (p = 0.47, η_p_^2^ = 0.04), nor significant interactions (ps > 0.25; Fig. 2).

**Figure 2.**
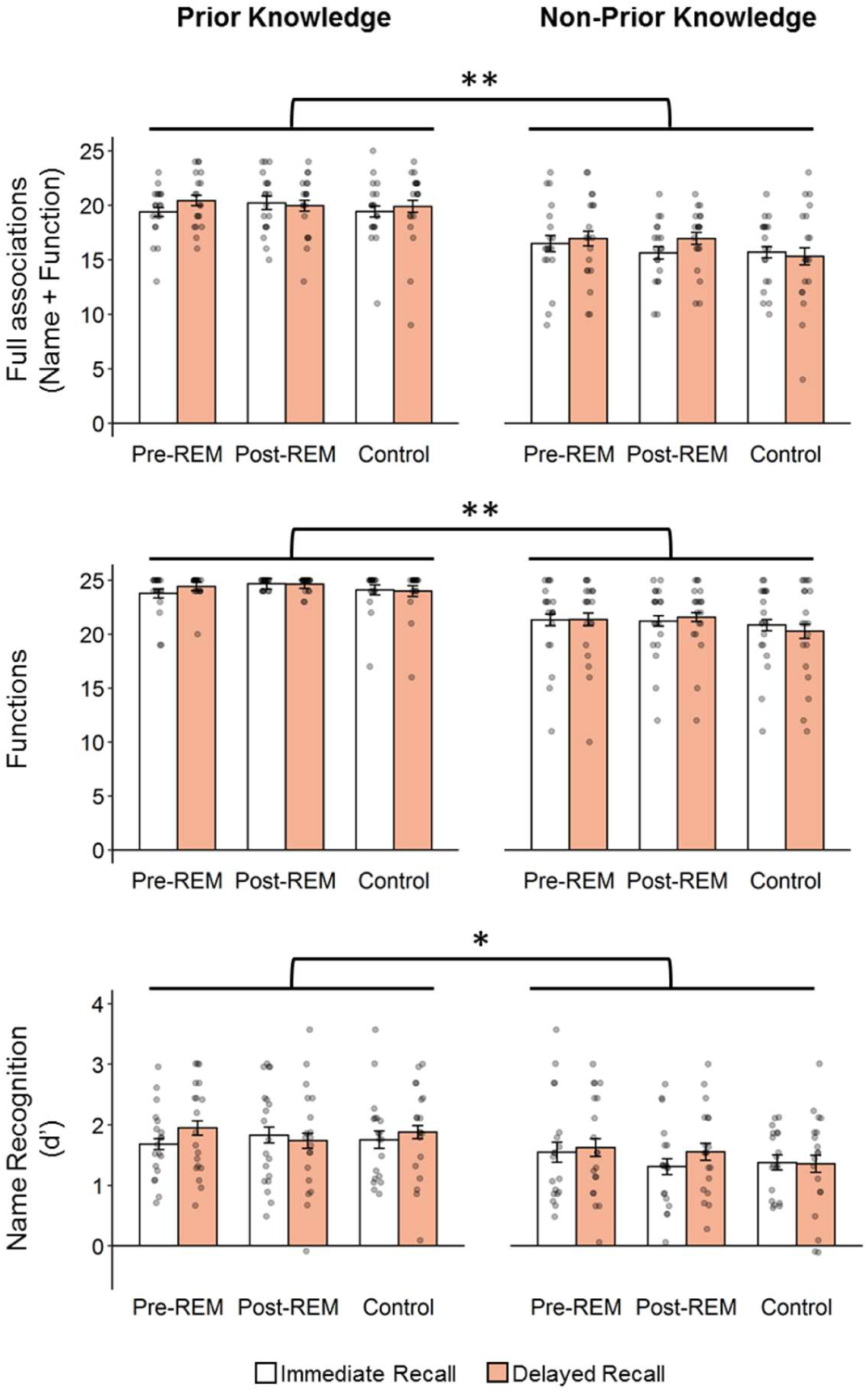
Overall performance (all items irrespective of cueing). Prior known associations are better remembered than non-prior known associations regardless of session (immediate vs. delayed recall), and condition (Pre-REM TMR, Post-REM TMR or Control), in all types of memory retrieval, i.e., fully remembered associations (top), functions (middle), and name recognition (bottom). Data are shown as mean ± SEM; ** p < 0.0001, * p < 0.001.

### Cueing effects on memory performance

We then compared the relative change in memory performance after TMR experimental sessions. The change for fully remembered associations at immediate recall, expressed as percentage change, did not differ between Pre-REM and Post-REM TMR conditions (p = 0.13, η_p_^2^ = 0.12), between cued and uncued subsets (p = 0.39, η_p_^2^ = 0.04), nor between prior and non-prior knowledge (p = 0.13, η_p_^2^ = 0.12). There were also no significant interactions (ps > 0.18) (Fig. 3a). Similarly, change in function recall did not differ significantly between TMR (p = 0.91, η_p_^2^ = 0.001), cueing (p = 0.2, η_p_^2^ = 0.09), and knowledge (p = 0.24, η_p_^2^ = 0.08) conditions, and no significant interactions emerged (all ps > 0.33) (Fig. 3a). In contrast, analysis of cueing effects on the relative change in name recognition accuracy (i.e., d’ change = delayed recall d’ – immediate recall d’) revealed a significant main effect of TMR condition, with better performance in the Pre-REM TMR condition as compared to the Post-REM TMR condition (F_1,18_ = 4.51, p = 0.048, η_p_^2^ = 0.2), regardless of cueing (p = 0.36, η_p_^2^ = 0.05) or knowledge (p = 0.49, η_p_^2^ = 0.03). There were no significant interaction effects (ps > 0.28) (Fig. 3b).

**Figure 3.**
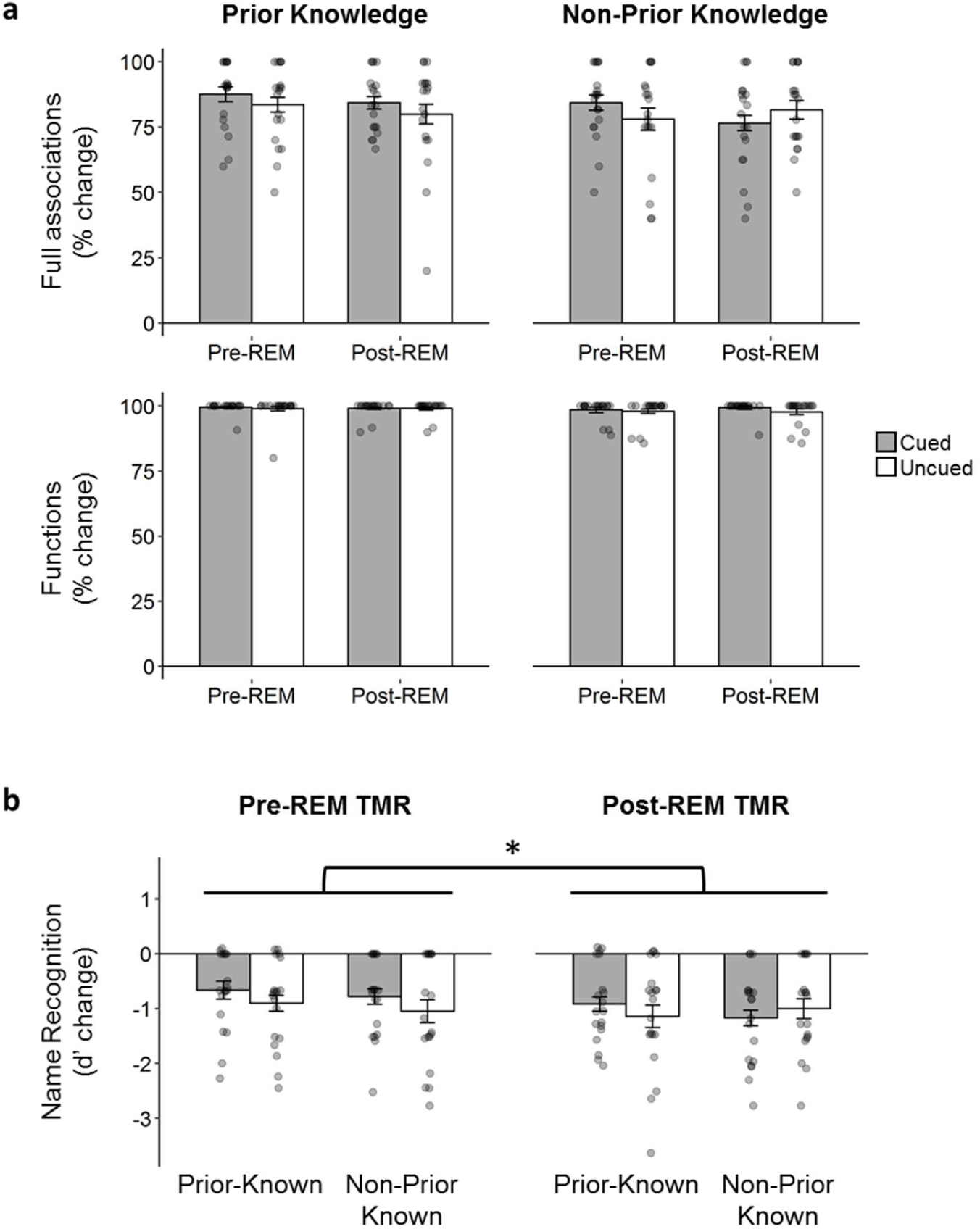
Cueing effects on correctly recalled items at immediate recall. **a** No significant differences in the relative change in memory performance for full associations and function recall across TMR (Pre-REM vs. Post-REM), cueing (cued vs. uncued words) and knowledge (prior vs. non-prior known). **b** Relative change in name recognition is significantly better in the Pre-REM TMR than in the Post-REM TMR condition, regardless of cueing or type of knowledge. Data are shown as mean ± SEM; * p < 0.05.

### Cueing effects on name recognition based on confidence ratings

To further understand the effect of TMR on name recognition, individual ROC curves were obtained according to confidence ratings provided by the participants in each condition at the delayed recall test, and their respective AUCs were calculated and compared. ROC curves averaged across participants per condition are shown (Fig. 4). Comparison of the ROC AUCs, showing the participants’ ability to discriminate correct from incorrect object names, revealed no main effects of TMR condition (p = 0.24, η_p_^2^ = 0.08), cueing (p = 0.7, η_p_^2^ = 0.01), or knowledge (p = 0.11, η_p_^2^ = 0.14). The condition by cueing interaction was significant (F = 5.39, p = 0.03, η_p_^2^ = 0.23), showing better name discrimination for cued associations in the Pre-REM TMR condition as compared to cued associations in the Post-REM TMR condition (t_18_ = 2.3, p = 0.03, d = 0.53) (Fig. 5). However, the difference between cued and uncued associations did not reach significance in the Pre-REM TMR condition (t_18_ = 1.67, p = 0.11, d = 0.38), nor between cued and uncued associations during Post-REM TMR (t_18_ = −0.98, p = 0.34, d = −0.23). No further significant interactions between TMR condition, cueing and knowledge factors were found (ps > 0.54).

**Figure 4.**
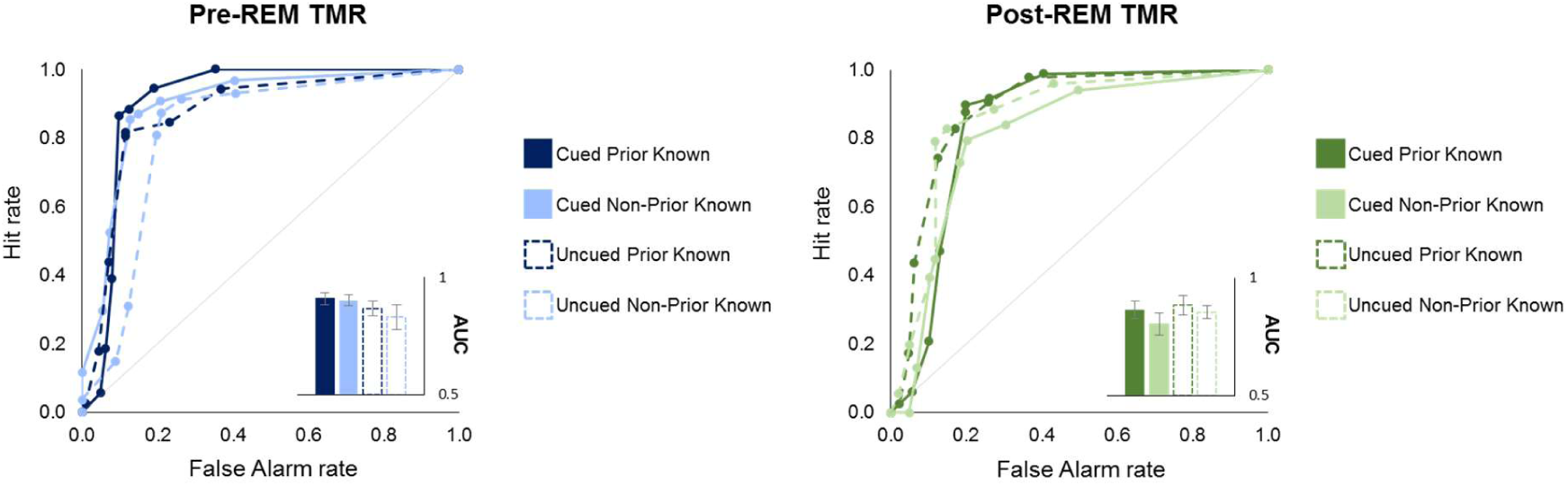
Cueing effects on confidence ratings for name recognition. ROC functions are computed for each cueing (cued, uncued words), and type of knowledge (prior known, non-prior known) condition for both Pre-REM (left) and Post-REM TMR conditions (right). Inset plots show the corresponding ROC AUCs. Data are shown as mean ± SEM.

**Figure 5.**
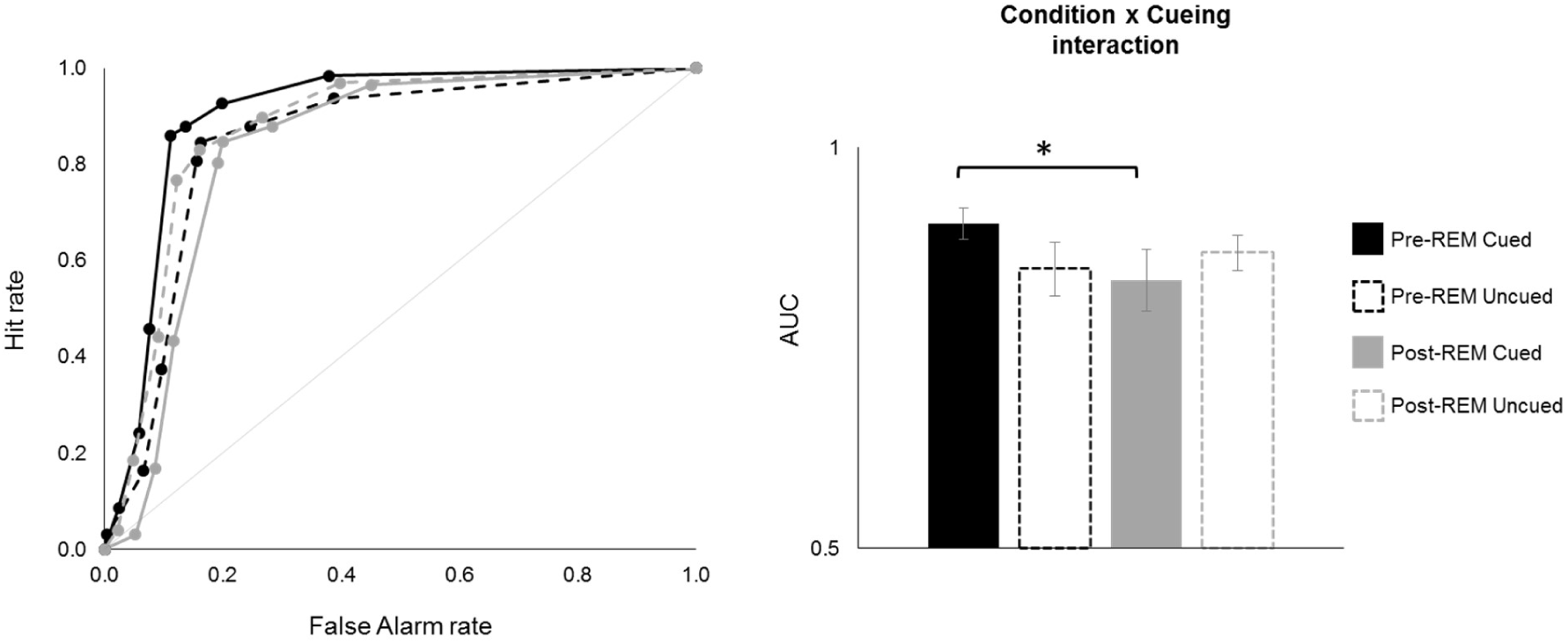
Condition by cueing interaction effects on confidence ratings for name recognition. ROC functions for cueing and TMR conditions were collapsed across types of knowledge (left). The ROC AUCs for cued associations in the Pre-REM TMR condition are significantly larger than ROC AUCs for cued items in the Post-REM TMR condition (right), suggesting an advantage for NREM-TMR cued associations when followed by REM sleep. Data are shown as mean ± SEM; * p < 0.05.

### Time-frequency analysis of cue-related activity

Changes in spectral power in response to cue presentation of subsequently remembered associations were compared across TMR conditions and types of knowledge. Comparison of TFRs between Pre- and Post-REM TMR conditions revealed a significant increase in sigma power during Pre-REM TMR, which ranged between 11-16 Hz, lasting approximately from 1.4 – 4.1 sec (p = 0.02, cluster-based permutation test; Fig. 6, Fig. S2) For a detailed characterisation of the cluster over each group of electrodes see Supplementary Fig. S3. There were no significant differences in relative power changes between prior and non-prior knowledge types of cues (ps > 0.15). Comparison of the mean difference between responses to prior known versus non-prior known cues in each TMR condition did not reveal significant interaction effects between these factors (ps > 0.4).

**Figure 6.**
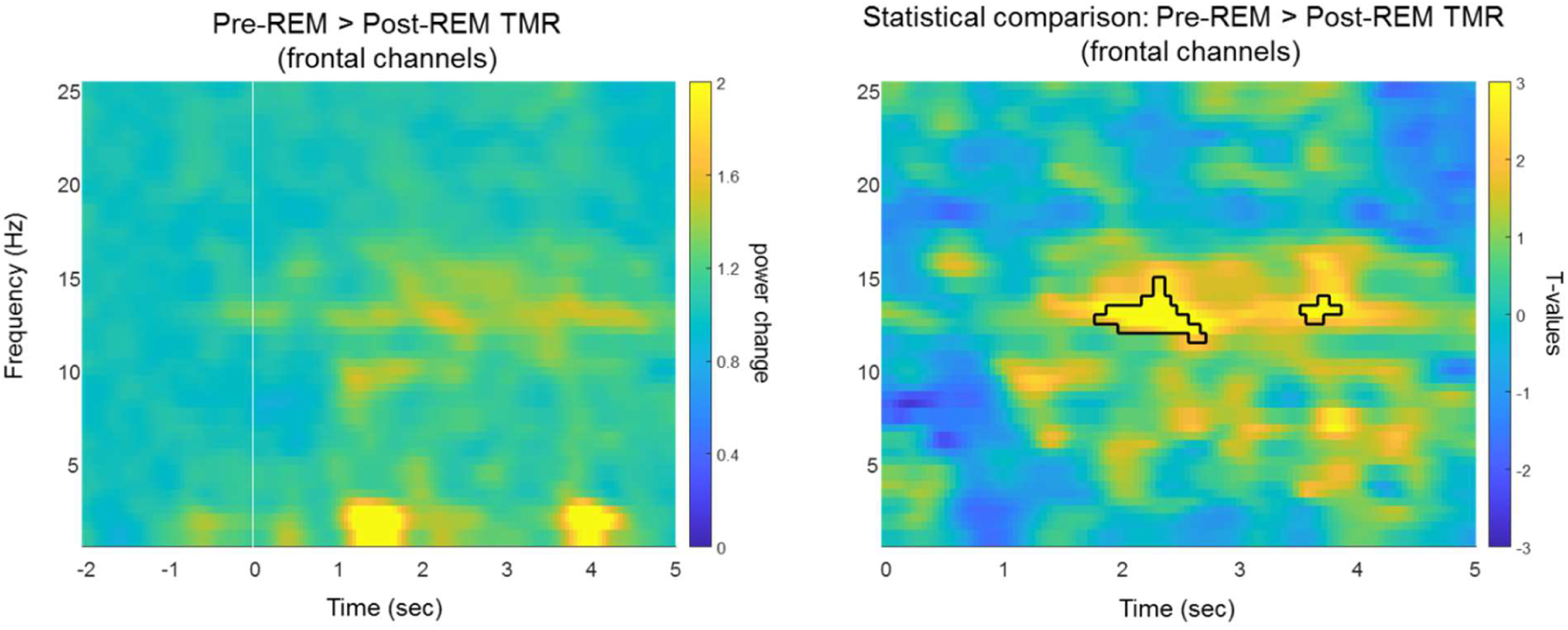
Time-frequency power changes during TMR over a representative cluster of electrodes. Left panel shows the spectral power contrast between Pre-REM and Post-REM TMR conditions over frontal channels in response to subsequently remembered cues (cue onset = 0 s). Right panel shows a cluster of activity in time and frequency, averaged over frontal channels, highlighting significant differences between Pre-REM and Post-REM TMR conditions (p = 0.02). Responses to cues are collapsed across prior and non-prior types of knowledge.

### Analysis of sigma power after cue presentation

To further investigate the cue-related changes in sigma activity and the behavioural outcomes of TMR, we analysed the correlation between sigma power (11-16 Hz) in response to cues within a time window defined by the start and end time of the identified cluster at each of the grouped electrode locations (see Supplementary Information Fig. S3), and the TMR benefit index for each condition and type of knowledge. Results disclosed a trend for a positive correlation between sigma power changes over frontopolar (r = 0.35, p = 0.079) and frontal (r = 0.38, p = 0.049) derivations and the TMR benefit for prior known associations cued in the Pre-REM TMR condition (Fig. 7a). A similar pattern of correlational trends emerged for the Post-REM TMR condition where sigma power over frontal (r = 0.36, p = 0.067), central (r = 0.50, p = 0.011) and parietal (r = 0.47, p = 0.021) channels correlated positively with the TMR benefit for prior known associations. However, these correlations did not survive FDR correction for multiple comparisons, and comparisons are thus reported for the sake of completeness (see Supplementary Information Table S1). No correlations between the change in sigma power after cue presentation and the TMR benefit for non-prior known associations were found in neither of the TMR conditions (ps > 0.24). Comparison between prior and non-prior known correlations did not reveal significant differences in the Pre-REM TMR condition (ps > 0.11) but for a difference at central channels (Z = 2.06; p = 0.04) in the Post-REM TMR condition (Fig. 7a). Significant differences between correlations were not revealed in none of the other locations (ps > 0.06; see Supplementary Information Table S3 for all correlation comparisons).

**Figure 7.**
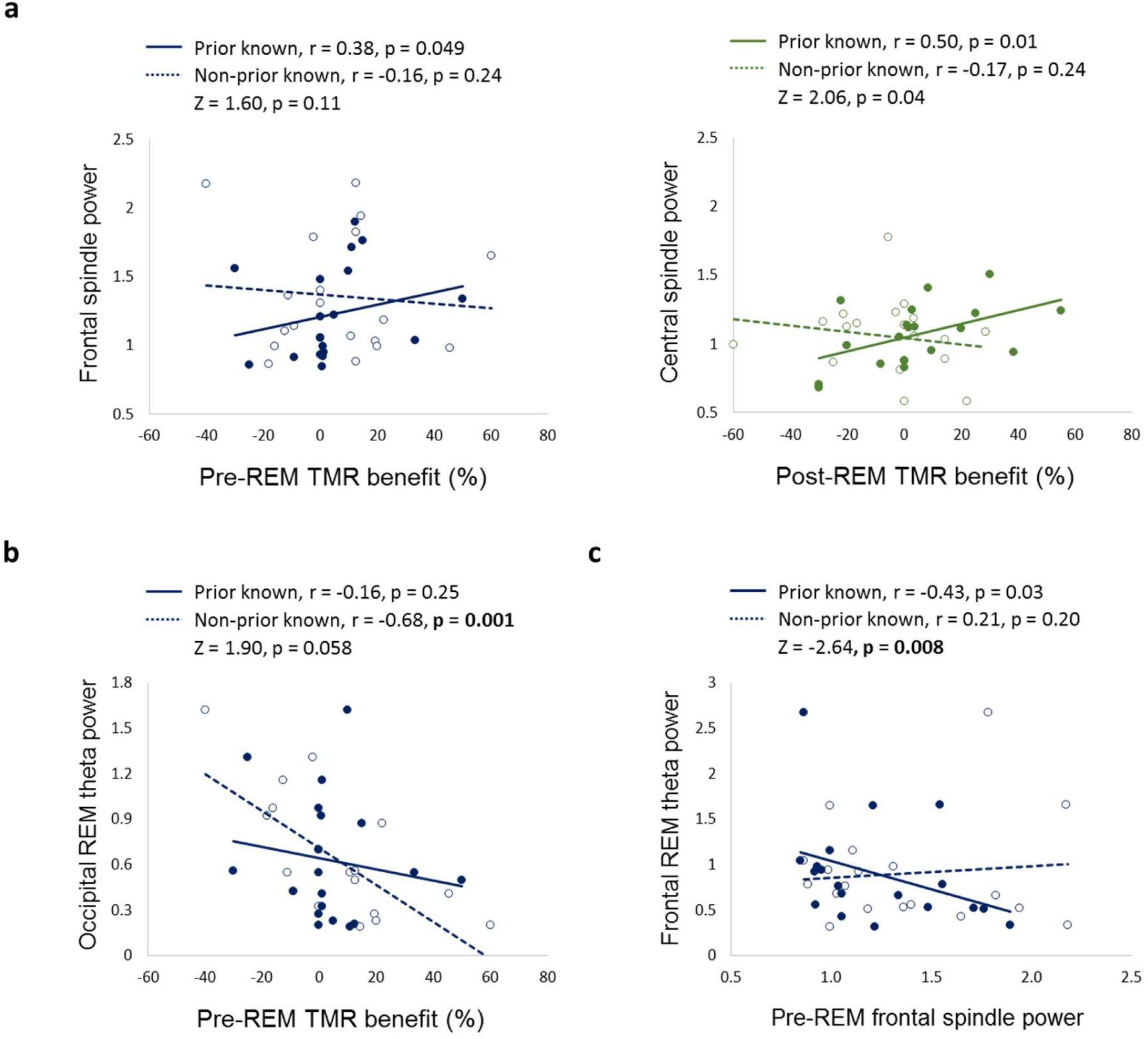
Correlation analyses results showing the relation between cue-evoked sigma activity, subsequent REM theta power and the TMR benefit index across conditions. **a** A positive association between evoked spindle power and the TMR memory benefit for prior known associations emerged in both Pre-REM and Post-REM TMR conditions. This was significantly different from the non-prior known correlation in the Post-REM TMR condition. **b** The Pre-REM TMR benefit index for non-prior known associations negatively correlated with subsequent REM theta power over posterior regions. **c** Sigma power evoked by reactivation of prior known associations negatively correlated with subsequent REM theta power over frontal regions. This correlation significantly differed from the non-prior known reactivation condition. Significant p-values after FDR correction are printed in bold.

### Analysis of REM sleep theta activity after NREM-TMR

Given the role of REM sleep theta activity in memory consolidation processes (Boyce et al. 2016, 2017), we investigated REM sleep theta power following Pre-REM TMR. We ran correlation analyses between REM sleep theta power (4-8 Hz) and the TMR benefit index of both, prior known and non-prior known associations. We did not find significant correlations between theta power and the TMR benefit for prior known associations (Table 2). Interestingly, significant negative correlations emerged between theta power and the TMR benefit index for non-prior known associations over all electrode groups, with higher correlations (r > - 0.53) over posterior regions (Table 2). It is worth noting that correlations between the Pre-REM TMR benefit for prior known associations showed trends towards a negative association with subsequent REM theta power as well (Table 2). Comparison between non-prior and prior known associations revealed a trend towards significance between correlations in the occipital electrode cluster (Z = 1.89, p-value = 0.058; Fig. 7b). No other correlations differed between the rest of locations (*ps* > 0.16).

**Table 2.** Correlations between Pre-REM TMR benefit index and subsequent REM theta power (4-8 Hz).

|  | Prior known |  | Non-prior known |  | Correlation comparison<br>(Steiger's Z-test) |  |
| --- | --- | --- | --- | --- | --- | --- |
|  | r | p | r | p | Z | p |
| Frontopolar | -0.37 | 0.055 | -0.48 | **0.014** | 0.38 | 0.71 |
| Frontal | -0.38 | 0.046 | -0.49 | **0.014** | 0.39 | 0.70 |
| Central | -0.32 | 0.086 | -0.61 | **0.003** | 1.06 | 0.29 |
| Temporal | -0.31 | 0.103 | -0.53 | **0.007** | 0.79 | 0.43 |
| Parietal | -0.33 | 0.071 | -0.67 | **0.001** | 1.31 | 0.19 |
| Occipital | -0.16 | 0.251 | -0.68 | **0.001** | 1.90 | 0.06 |

Significant p-values after FDR correction are printed in bold.

Finally, we conducted an analysis on the association between sigma power (11-16 Hz) in response to both types of cues in the Pre-REM TMR condition, and subsequent REM theta power. There was a significant negative correlation between sigma power changes in response to prior known cues and subsequent REM sleep theta power over frontal electrodes (r = - 0.43; p = 0.03), however this correlation did not survive FDR correction (Fig. 7c). Central (p = 0.049) and frontopolar (p = 0.079) electrode clusters showed similar trends (see Supplementary Information Table S2). As for non-prior known cue-evoked spindle activity, no correlations with subsequent REM theta activity were found. Further comparison between correlations revealed significant differences between prior and non-prior knowledge cueing over frontopolar (Z = 2.55; p = 0.01), and frontal electrodes (Z = 2.64; p = 0.008) (Fig. 7c). No other correlations significantly differed over other electrode locations (*ps* > 0.09; see Supplementary Information Table S2 for all correlation comparisons).

## Discussion

The present study aimed at investigating whether a naturally following episode of REM sleep is necessary for prior NREM TMR to exert a beneficial effect on the consolidation of novel word associations. In addition, we investigated whether this potential benefit interacts with the level of prior knowledge involved in the newly learned associations. We performed a within-subject study in which TMR was delivered in the second half of the night either during NREM sleep followed by REM sleep, or during NREM after the last episode of REM sleep. Targeted memories included both prior known and non-prior known associations (i.e., pseudowords associated with familiar or novel object pictures). In line with sequential accounts of sleep for memory consolidation (Giuditta et al. 1995; Ambrosini and Giuditta 2001) and TMR studies pointing to a benefit of REM sleep after NREM-TMR (Batterink et al. 2017; Tamminen et al. 2017), we predicted that NREM-TMR would be beneficial for memory performance when followed by a period of REM sleep (Pre-REM TMR) as compared to a period of NREM-TMR not completed by REM sleep. In particular, we hypothesized that cued associations involving a higher degree of prior knowledge would benefit to a greater extent from the Pre-REM TMR condition, due to their higher connections with pre-existing knowledge networks and to the effect of REM sleep in integrating those associations (Walker and Stickgold 2010). On the other hand, we hypothesized that cueing memories during NREM sleep without a following period of REM sleep (Post-REM TMR) might lead to destabilization of memory traces, and consequently a decrease in memory performance for cued associations, particularly in those involving less connections with pre-existing networks.

Our results only partially support these predictions. At the behavioral level, we compared the relative change in memory performance for cued vs uncued associations that were fully remembered at immediate recall. Contrary to our predictions, there were no differences between TMR (Pre-REM vs. Post-REM), cueing (cued vs. uncued), and knowledge (prior known vs. non-prior known) for full association and function recall. However, accuracy in name recognition was significantly better in the Pre-REM TMR than the Post-REM TMR condition, regardless of cueing or knowledge type — i.e., a selective benefit for cued associations was still not evident. A further analysis conducted on confidence ratings for name recognition showed a significant interaction between TMR and cueing factors, indicating significantly larger ROC AUCs for cued words in the Pre-REM TMR condition relative to cued words in the Post-REM TMR, regardless of knowledge. Still, the expected difference between cued and uncued associations within each of the TMR conditions was not revealed. On the other hand, prior knowledge did not seem to benefit to a greater extent from Pre-REM TMR, or from the TMR procedure in general, at least from a behavioural standpoint.

At the neural level, we observed increased sigma power (reflecting sleep spindles) in the Pre-REM TMR condition for later remembered cues compared to the Post-REM TMR condition. Unlike a prior study (Groch et al. 2017), we found no differences in power changes between prior and non-prior known associations or interactions between TMR conditions and type of knowledge. Nevertheless, correlation analyses between evoked sigma power and the TMR benefit index for prior known associations revealed positive associations in both TMR conditions.

Finally, we examined whether subsequent REM theta activity played a role in supporting memory reactivation processes induced during prior NREM-TMR. We found significant negative correlations between REM sleep theta activity and the TMR benefit index for non-prior known associations in the Pre-REM TMR condition. Correlation trends were also observed for prior known items. Descriptively, negative correlations between theta power and the TMR benefit for non-prior associations were more pronounced over posterior regions. Furthermore, we found a negative association between evoked sigma activity for prior known cues during Pre-REM TMR and subsequent REM theta power in frontopolar and frontal regions which significantly differed from the correlation between evoked sigma for non-prior known associations and subsequent REM theta, suggesting that potential differences in cue-evoked sigma power may become relevant for memory processes during subsequent REM sleep.

### REM sleep after NREM TMR and novel word recall

Looking at overall performance changes, we found an advantage for prior known associations across recall sessions in TMR and control conditions, consistent with well-established schema-based learning effects (Tse et al. 2007; Van Kesteren et al. 2012). Additionally, prior research suggested that sleep facilitates memory consolidation for familiar or schema-conformant information (Durrant et al. 2015; Hennies et al. 2016). However, sleep-mediated memory benefits may also extend to more difficult, unfamiliar associations when additional practice is provided during the encoding phase (Cordi et al. 2023). In our study, we did not find an overnight advantage for prior over non-prior known associations — but a main effect of knowledge. By design, it was expected that during the learning and practice rounds, participants would initially retrieve a higher number of prior known associations, as the 60% performance criterion we established did not distinguish between types of associations. In average, participants required at least 2 learning and practice rounds to achieve this criterion, resulting in more repetitions of non-prior known associations (as successfully retrieved associations were dropped from the subsequent practice round). It is possible that weaker encoding of non-prior known associations was compensated through re-exposure and practice, leading to no differences in overnight performance for both types of knowledge — with generally better memory for prior known associations. Furthermore, the lack of TMR benefits in overall performance could be explained by the effect of multiple learning and practice rounds in strengthening memories before sleep, as it has been shown that TMR favours weaker representations (Cairney et al. 2016), and its beneficial effects may be reduced by prior testing (Joensen et al. 2022). To better understand the effects of sleep and cueing on memory with respect to prior knowledge, future studies should ensure similar encoding strengths by setting separate intermediate learning criteria for prior and non-prior knowledge items, and carefully address how these memories are established during learning, for example by taking into account potential differences between re-studying and re-testing items with varying degrees of prior knowledge (Antony et al. 2017).

To investigate cueing effects, we compared the relative change in memory performance across TMR, cueing and knowledge conditions. We found no significant differences across these factors for full associations or function recall. Name recognition was better in the Pre-REM TMR condition — yet, no significant differences between cued and uncued associations nor between types of knowledge emerged. Further analysis of name discrimination based on confidence ratings, as shown by ROC AUCs, revealed that the advantage observed in the Pre-REM TMR condition was specific to the cued associations. This advantage was regardless of prior knowledge, as indicated by a significant interaction between TMR and cueing. Specifically, Pre-REM cued associations, which exhibited the highest ROC AUCs, significantly differed from Post-REM TMR cued associations, which showed the lowest ROC AUCs. However, the advantage for cued versus uncued items in the Pre-REM TMR condition did not reach significance. One interpretation for these results could partially align with the prediction that additional REM sleep may take part in memory consolidation processes initiated by NREM TMR, while reactivated memories without subsequent REM sleep may be destabilized. This pattern of results is in line with two previous studies showing no cueing effects of TMR over an afternoon nap for pseudoword-picture learning (Batterink et al. 2017) and lexical competition (Tamminen et al. 2017), but an influence of REM sleep duration on the effect of TMR. While participants who spent more time in REM sleep exhibited increased TMR memory benefits, participants who spent little or no time in REM sleep showed a detrimental effect of cueing (Batterink et al. 2017; Tamminen et al. 2017).

An alternative interpretation, and a potential limitation inherent to our design, is that the lack of selective enhancement for cued versus uncued associations across memory measures, and the rather weak effect of Pre-REM TMR in name recognition measures (i.e., main effect of change in d’ and higher Pre-REM cued versus Post-REM cued ROC AUCs) could be explained by the timing of TMR delivery. Specifically, after immediate recall, participants were allowed to sleep uninterruptedly until the last two cycles of the night, that we targeted due to the presence of REM episodes that are longer and richer in phasic events (Simor et al. 2020). It is possible that spontaneous reactivations occurring during the first sleep cycles, characterized by a higher presence of SWS, may have initiated the consolidation of newly learned material, eventually leading to reduced effectiveness of TMR interventions conducted later in the night. Support from this notion comes from a rodent study showing that the presentation of task-related cues successfully biased the spontaneous occurrence of neuronal replay events only during the first, but not the second half of the sleep session (consisting of one or more sleep/wake cycles) (Bendor and Wilson 2012). Moreover, and in contrast with our results, other vocabulary learning studies have shown that delivering TMR during the first episodes of NREM sleep led to significant selective cueing memory enhancement in foreign language word pairs (Schreiner and Rasch 2015; Schreiner et al. 2015) and pseudowords paired with familiar objects (Groch et al. 2017). To the best of our knowledge, only a few studies initiated TMR during the second half of the night (e.g., Sterpenich et al. 2014; Rihm and Rasch 2015; Laventure et al. 2016). In one of these studies (Rihm and Rasch 2015), to avoid the spontaneous reactivation of task-related memories during SWS, participants were allowed to sleep during the first hours of the night, after which they were awoken to perform an aversive conditioning learning task and went back to sleep during which TMR was performed. Even though they found an effect of TMR in overall arousal ratings, no effect of NREM stage 2 TMR was observed on memory performance. Together with our current findings, and as shown by numerous studies where TMR took place in the initial episodes of sleep (see Hu et al. 2020 for a review), TMR is likely to be most effective when performed during the first cycles of sleep (Simor et al. 2023). Additionally, we cannot entirely rule out potential neuromodulatory or circadian influences on cueing effects. For instance, blocking the early morning cortisol rise after reactivating consolidated memories has been shown to enhance memory performance (Antypa et al. 2021). On the other hand, we find it unlikely that factors like arousal or limited cue repetitions due to lighter sleep negatively impacted Post-REM TMR. Cue repetition and the proportion of NREM sleep stages 2 and 3 were similar during the stimulation period of both TMR nights, and the proportion of time spent awake during this period was actually higher for Pre-REM TMR.

### Neural responses to NREM TMR, subsequent REM sleep, and their association with TMR effects on prior knowledge

Sigma power in response to later-remembered cues was significantly higher in the Pre-REM TMR compared to the Post-REM TMR condition as shown by the TFR analysis. No further differences between types of knowledge or interaction effects were found. This result is in line with previous studies showing increases in sigma activity after cue presentation (Schreiner et al. 2015; Cairney et al. 2017; Farthouat et al. 2017). Despite no observed differences in evoked power between prior known and non-prior known cues, we found that the TMR benefit index for prior known associations positively correlated with evoked sigma power at fronto-central and parietal regions in Pre- and Post-REM TMR conditions respectively, whereas such pattern of associations was not observed for non-prior known associations in neither of the TMR conditions. These results are in line with previous work showing that increased spindle activity elicited by cueing of prior known associations is predictive of post-sleep TMR benefits for memory performance (Groch et al. 2017), and with the proposal that spindle activity mediates the integration of novel information into pre-existing knowledge networks (Tamminen et al. 2010, 2013; Hennies et al. 2016). However, these results should be considered with caution, and independently confirmed, as significant correlations between spindle activity and the TMR benefit for prior known associations did not survive correction for multiple comparisons.

While circadian variations may account for the increased spindle power during Pre-REM TMR, independently of subsequent REM sleep, we found different associations between cue-evoked sigma activity, subsequent REM theta power, and the TMR benefit index, suggesting distinctive interactions between NREM and REM sleep for memory consolidation. This interplay was further influenced by the type of knowledge being reactivated. For instance, we found significant negative associations between REM sleep theta power and the Pre-REM TMR benefit index of non-prior known associations, and a similar pattern (although only at trend levels) for prior known associations. These results are surprising given the proposed role of REM theta activity for memory consolidation (Boyce et al. 2016, 2017). Notwithstanding, REM sleep was also found to be implicated in synaptic depotentiation, or forgetting (Poe 2017), with support for this notion coming from studies showing a role of REM sleep in selective pruning of synapses (Li et al. 2017), active forgetting of hippocampus-dependent memories (Izawa et al. 2019) and downscaling of hippocampal and neocortical excitability (Grosmark et al. 2012; Miyawaki and Diba 2016; Watson et al. 2016). In humans, studies found negative associations between REM sleep and perceptual (Strauss et al. 2022) and gross motor skills (Hoedlmoser et al. 2015). In our study, TMR benefits for both types of knowledge showed negative associations with REM theta, yet only the correlations with non-prior knowledge cueing were significant. We may speculate that while reactivation of prior knowledge associations may have promoted their stabilization and integration into pre-existing networks — as reflected by positive associations between spindle power and prior knowledge cueing benefits — reactivation of non-prior known associations involving less overlap with pre-existing networks may have even been weakened (Lewis and Durrant 2011). In a second step taking place during REM sleep, potential downscaling processes may have been more deleterious for non-prior known cued memories — as reflected by the strong negative associations between REM theta power and non-prior knowledge cueing benefits. The fact that these correlations were stronger in posterior areas is intriguing and deserves further investigation. One possibility is that the observed pattern of correlations was related to the kind of learned associations, as it has been shown that posterior temporal and parietal cortical areas are involved in the integration of visual representations of objects, phonological presentation of novel words, and related semantic information (Takashima et al. 2017). The explanation of these results are admittedly speculative, and should be substantiated by further research showing clear behavioural differences between prior and non-prior knowledge reactivated memories.

The hypothesis that NREM TMR and REM sleep act synergistically in the processing of reactivated memories was further supported by an additional analysis showing a negative correlation between spindle activity evoked by prior known cues at frontal regions and subsequent REM theta activity. These results are in agreement with previous studies showing a complex interplay between sequential NREM and REM sleep, and related oscillatory phenomena for memory processing. In a rare opportunity to directly test the role of ordered NREM-REM sequences in memory consolidation, Strauss et al. (2022) tested patients with narcolepsy — who may often start sleeping directly in REM — on a visual perceptual skills task before and after a one-cycle daytime nap comprising either a REM-NREM or NREM-REM sleep sequence. Even though performance was similar and remained stable after both types of naps, the authors found that spindle duration positively correlated with post-sleep performance only when NREM sleep was followed by REM sleep, and that REM sleep phasic events (rapid eye movements) negatively correlated with skill performance. In another human study using magnetic resonance spectroscopy (Tamaki et al. 2020), the excitatory and inhibitory (E/I) balance (i.e., an index of plasticity) was studied in visual areas during sleep following training on a visual skills task. The authors found that while NREM sleep promotes plasticity associated with post-sleep performance gains, training-related plasticity decreases in subsequent REM sleep promoted the stabilization of the trained skill. Interestingly, and similar to our pattern of results, they also found that NREM spindle and REM theta power positively correlated with performance gains and stabilization of the skill respectively, whereas REM theta power also negatively correlated with the E/I balance. As for declarative memory tasks, Blaskovich et al. (2017) showed that recall of relevant words in a directed forgetting paradigm was better after an afternoon nap only when participants reached REM sleep. Moreover, spindle activity and subsequent REM duration positively correlated with recall of relevant words, suggesting that besides a sequential role of NREM spindle activity and REM sleep for consolidation processes, memories may be selected and differentially processed depending on their future relevance. Further evidence has shown that the interplay between sequential NREM and REM sleep oscillatory patterns may also impact the homeostatic regulation of neuronal activity. For instance, down-regulation of neocortical and hippocampal firing rates during sleep was observed to decrease over interleaving bouts of REM sleep (Grosmark et al. 2012; Miyawaki and Diba 2016), and this decrease in firing was predicted by spindles and SWRs only when NREM episodes were separated by REM sleep. Moreover, spindle rates from preceding NREM epochs predicted subsequent REM sleep theta power (Miyawaki and Diba 2016).

Although our findings did not fully support a beneficial effect of REM sleep after NREM TMR for the consolidation of reactivated memories, a complex relationship between memory reactivation processes triggered by TMR during prior NREM and subsequent REM sleep seems to emerge. Whether the fate of reactivated memories depends not only on the sequence of sleep stages, and related oscillatory phenomena, but also on the type of associations that are reactivated requires further investigation.

## Acknowledgements

The authors would like to thank Flavia Chiovini and Katja Sandkühler for their assistance in data collection. R.S.O was supported by the Fonds de la Recherche Scientifique (F.R.S.-F.N.R.S., Aspirant Research Fellowship). The study was also partially supported by the F.N.R.S. and the Fonds Wetenschappelijk Onderzoek – Vlaanderen (F.W.O.) under the Excellence of Science (EOS) Project (MEMODYN, No. 30446199 to P.P.). P.S. was supported by the Hungarian National Research, Development, and Innovation Office Grant NKFI FK 142945, and by the Janos Bolyai scholarship of the Hungarian Academy of Sciences.

