## Supplementary Information for "Does targeted memory reactivation during NREM sleep require complementary REM sleep for memory consolidation?"

1. **Progression through practice rounds after the learning phase**

Prior to the immediate recall phase, participants had to remember at least 60% of the complete associations (i.e., recalling both name and function of the object). In order to achieve this criterion participants required in average 1.9 rounds of practice (Pre-REM TMR mean = 1.89 ± 0.46; Post-REM TMR = 1.89 ± 0.46; Control = 2.05 ± 0.52). At the descriptive level, more prior known associations were correctly recalled in the first round of practice (Fig. S1), while more non-prior knowledge associations were correctly recalled in the following round.

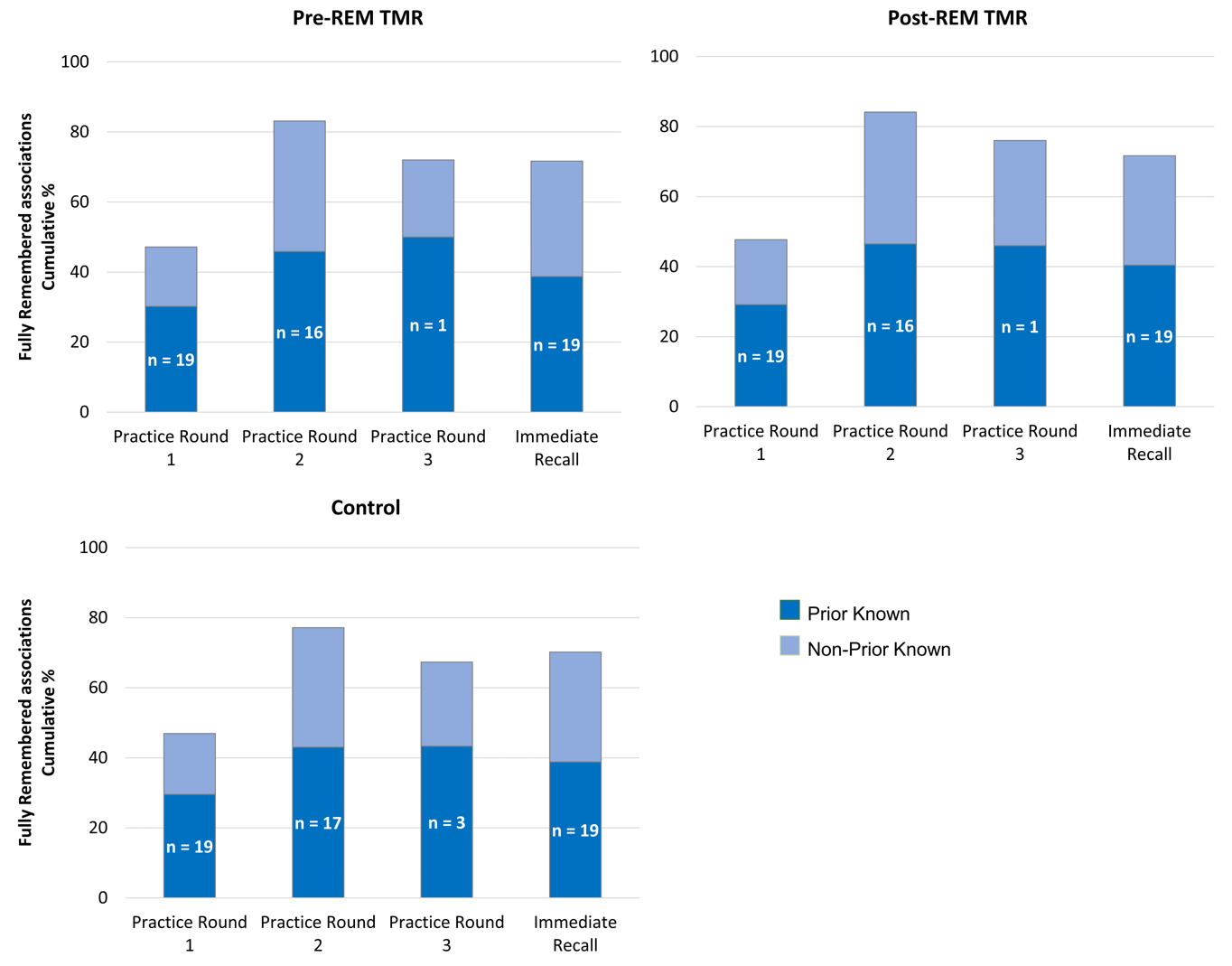

**Figure S1.** After the learning phase, participants required ~2 practice rounds to reach the 60% criterion in order to pass to the immediate recall test before sleep (performance in this latter test is shown in the last bar of each subplot). Bars show the cumulative percent of fully remembered associations for prior and non-prior known object associations across TMR and Control conditions. Number of participants per round is also indicated.

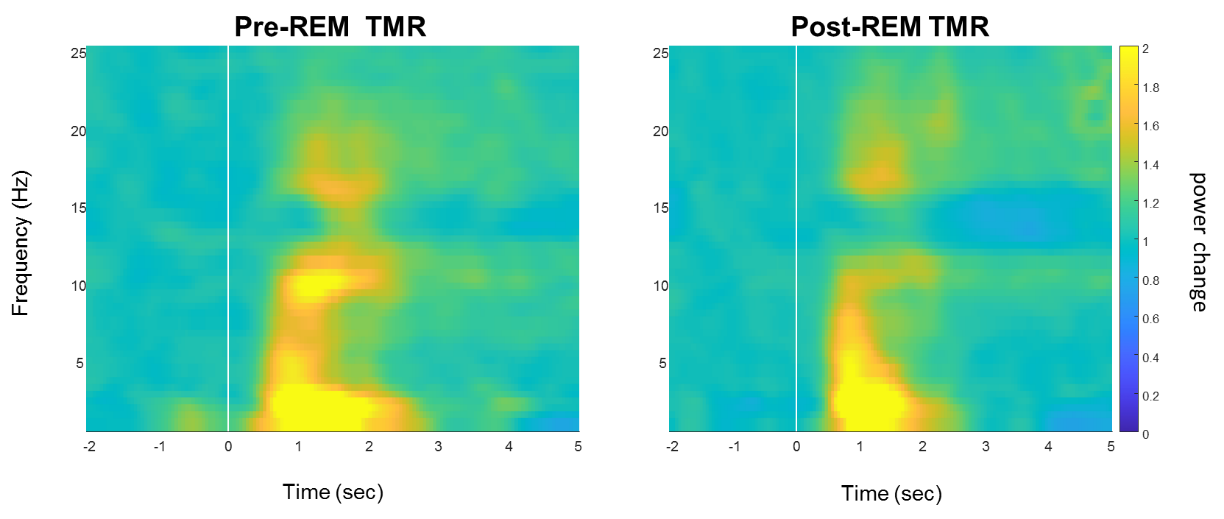

**Figure S2.** Time-frequency power changes after presentation of subsequently remembered cues are shown for Pre-REM and Post-REM TMR conditions (cue onset = 0 s). Responses to cues are collapsed across prior and non-prior types of knowledge.

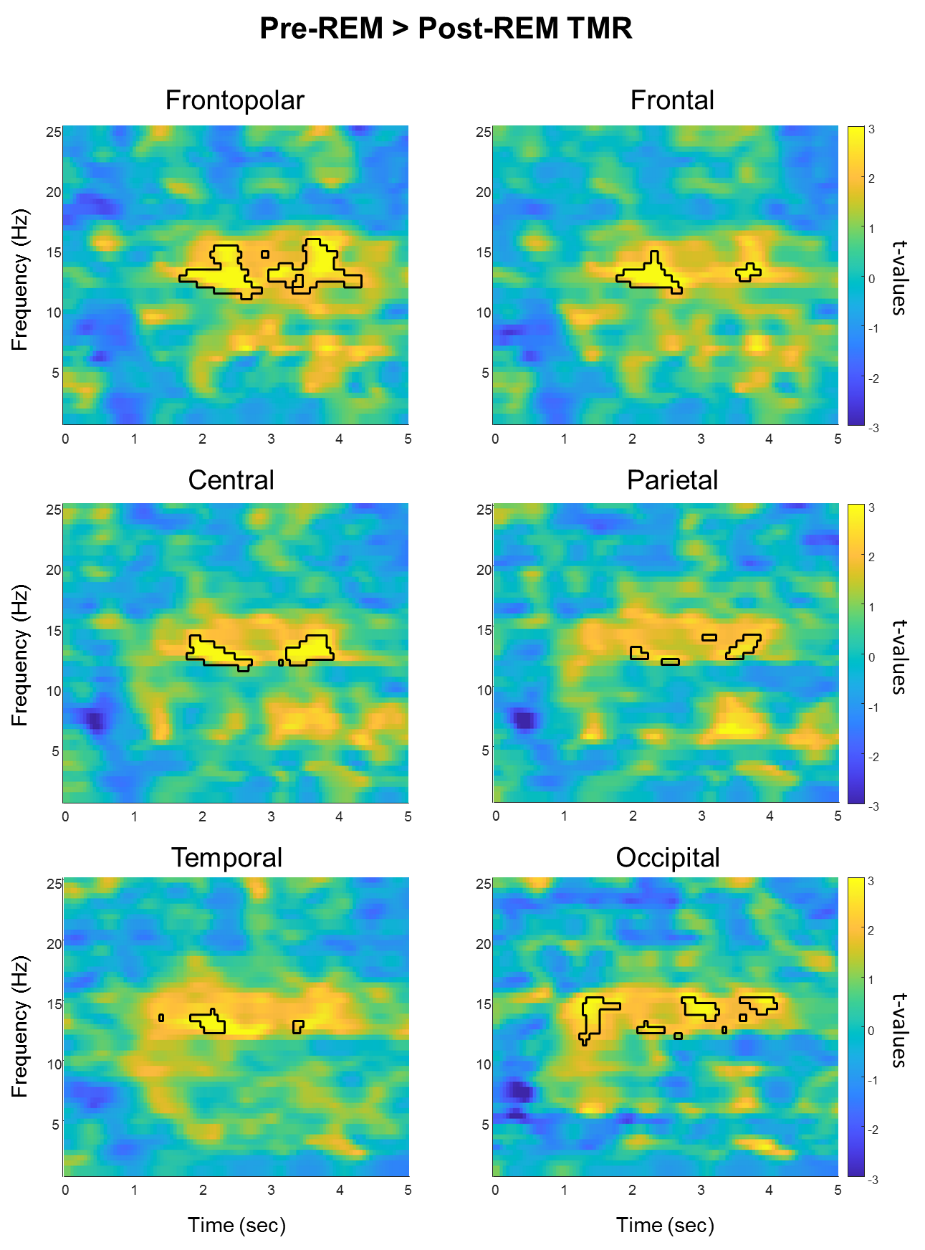

**Figure S3.** Significant time-frequency power differences emerged over all electrodes after cue-presentation in the Pre-REM TMR as compared to the Post-REM TMR condition. Panels show significant differences averaged over electrode locations (p = 0.02).

| **Table S1.** Correlations between TMR benefit index and spindle activity after cue presentation. | | | | | | |
| --- | --- | --- | --- | --- | --- | --- |
|  | Prior known | | Non-prior known | | Correlation comparison (Steiger’s Z-test) | |
|  | r | p | r | p | Z | p |
| **Pre-REM TMR** |  |  |  |  |  |  |
| Frontopolar | 0.35 | 0.079 | -0.13 | 0.304 | 1.41 | 0.16 |
| Frontal | 0.38 | 0.049 | -0.16 | 0.242 | 1.60 | 0.11 |
| Central | 0.23 | 0.160 | -0.09 | 0.386 | 0.94 | 0.35 |
| Temporal | 0.32 | 0.095 | -0.11 | 0.354 | 1.25 | 0.21 |
| Parietal | 0.15 | 0.274 | -0.04 | 0.469 | 0.56 | 0.57 |
| Occipital | 0.04 | 0.450 | 0.06 | 0.387 | -0.06 | 0.95 |
| **Post-REM TMR** |  |  |  |  |  |  |
| Frontopolar | 0.23 | 0.170 | 0.07 | 0.384 | 0.46 | 0.65 |
| Frontal | 0.36 | 0.067 | -0.03 | 0.456 | 1.16 | 0.25 |
| Central | 0.50 | 0.011 | -0.17 | 0.240 | 2.06 | 0.04 |
| Temporal | 0.26 | 0.149 | 0.06 | 0.414 | 0.57 | 0.57 |
| Parietal | 0.47 | 0.021 | -0.12 | 0.300 | 1.87 | 0.06 |
| Occipital | 0.03 | 0.433 | -0.06 | 0.363 | 0.28 | 0.78 |

Pearson correlations between the TMR benefit index for both conditions and evoked spindle power after cue onset. Significant correlation p-values after FDR correction are printed in bold. P-values did not survive FDR correction for multiple comparisons.

| **Table S2.** Correlations between spindle activity (11-16 Hz) evoked by Pre-REM TMR cueing and subsequent REM theta power (4-8 Hz). | | | | | | |
| --- | --- | --- | --- | --- | --- | --- |
|  | Prior known | | Non-prior known | | Correlation comparison (Steiger’s Z-test) | |
|  | r | p | r | p | Z | p |
| Frontopolar | -0.35 | 0.07 | 0.32 | 0.10 | -2.55 | **0.01** |
| Frontal | -0.43 | 0.03 | 0.21 | 0.20 | -2.64 | **0.008** |
| Central | -0.37 | 0.05 | 0.02 | 0.47 | -1.54 | 0.12 |
| Temporal | -0.20 | 0.21 | 0.22 | 0.18 | -1.66 | 0.10 |
| Parietal | -0.24 | 0.15 | -0.14 | 0.31 | -0.41 | 0.68 |
| Occipital | 0.04 | 0.46 | -0.21 | 0.20 | 1.18 | 0.24 |

Pearson correlations between spindle activity evoked by Pre-REM TMR cues and subsequent REM sleep theta power. Significant correlation p-values after FDR correction are printed in bold.
